# Amplitude of high frequency oscillations as a biomarker of the seizure onset zone

**DOI:** 10.1101/2020.06.28.176222

**Authors:** Krit Charupanit, Indranil Sen-Gupta, Jack J Lin, Beth A Lopour

## Abstract

**Objective:** Studies of high frequency oscillations (HFOs) in epilepsy have primarily tested the HFO rate as a biomarker of the seizure onset zone (SOZ), but the rate varies over time and is not robust for all individual subjects. As an alternative, we tested the performance of HFO amplitude as a potential SOZ biomarker using two automated detection algorithms.

**Method:** HFOs were detected in intracranial electroencephalogram (iEEG) from 11 patients using a machine learning algorithm and a standard amplitude-based algorithm. For each detector, SOZ and non-SOZ channels were classified using the rate and amplitude of high frequency events, and performance was compared using receiver operating characteristic curves.

**Results:** The amplitude of detected events was significantly higher in SOZ. Across subjects, amplitude more accurately classified SOZ/non-SOZ than rate (higher values of area under the ROC curve and sensitivity, and lower false positive rates). Moreover, amplitude was more consistent across segments of data, indicated by lower coefficient of variation.

**Conclusion:** As an SOZ biomarker, HFO amplitude offers advantages over HFO rate: it exhibits higher classification accuracy, more consistency over time, and robustness to parameter changes.

**Significance:** This biomarker has the potential to increase the generalizability of HFOs and facilitate clinical implementation as a tool for SOZ localization.

## 1. Introduction

High frequency oscillations (HFOs), defined as distinguishable spontaneous electrographic patterns consisting of at least four cycles in the frequency range of 80–500 Hz, are a promising biomarker for the seizure onset zone (SOZ) [1–8]. HFOs are commonly categorized as ripples, ranging from 80 to 250 Hz [9,10], and fast ripples, ranging from 250 to 500 Hz [11,12]. Both ripples and fast ripples occur more frequently in the SOZ than outside the SOZ in human intracranial electroencephalogram (iEEG) recordings [1–8]. Moreover, retrospective studies have reported that removal of brain regions with high occurrence of interictal HFOs is correlated to a positive outcome after epilepsy surgery [1,2,13–18]. HFOs may therefore be a valuable tool for localization of epileptogenic tissue to aid in surgical planning [19]. In addition, because HFO analysis is typically done on interictal iEEG data, this marker represents an opportunity to reduce the duration of the pre-surgical monitoring period [20,21].

The HFO rate (number of HFOs per minute per channel) has been almost exclusively used as the metric to test HFOs as a biomarker of the SOZ [1,2,4–8,22–27]. However, there are several limitations to this method. The mean interictal HFO rate varies across brain regions, for both epileptic and non-epileptic tissue [14,28,29], which complicates selection of a threshold for classification of pathological and physiological regions. The rate is also time-dependent, as it increases during sleep [8,30] and has different mean values in interictal, ictal, and preictal states [26,31,32]. Even when only interictal non-REM sleep data is used, the estimated location of the SOZ based on HFO rate can change over time [33]. Moreover, while it is understood that a high rate of HFOs may indicate epileptogenic tissue, the wide variability of the rate necessitates a patient-specific interpretation of this measurement. For example, the reported rates of HFOs in the SOZ vary more than an order of magnitude, from 1-2 HFOs per minute per channel [1,34] to greater than fifty per minute per channel [6,8,35]. This measurement is also confounded by the occurrence of physiological HFOs, which are associated with healthy cognitive processing [36]. The characteristics of physiological and pathological HFOs are not fully understood and may overlap with one another [37,38]. Lastly, the detection method and parameters can affect the measurement of HFO rate. If an energy threshold is used in the HFO detection procedure, the estimate of HFO rate will be a function of the threshold, with lower values producing higher rates. Similarly, a detector that is susceptible to false positive detections may suggest a higher HFO rate than a more specific detector applied to the same dataset.

As alternatives to rate, differences in other HFO features, such as amplitude, peak frequency, and duration have been reported [2,23,27,37]. In particular, several studies reported secondary results suggesting that HFO amplitude is significantly higher in SOZ than non-SOZ (nSOZ) channels for both ripples [22,23,26,27,39] and fast ripples [27,39]. However, while significant, these differences in amplitude were often small, and they were sometimes only seen in a subset of brain regions [14]. We hypothesized that HFO amplitude is higher in the SOZ, but the effect size may be reduced by the choice of detection methods and parameters. For example, if an energy-based threshold is used for detection, as the threshold increases, so will the estimated mean amplitude for both SOZ and nSOZ channels. Similarly, visual detection of HFOs will be biased toward the highest amplitude events, in both SOZ and nSOZ regions.

Therefore, we tested the performance of HFO amplitude as a biomarker of the SOZ by applying two different automatic detection algorithms to human iEEG with varying parameters. First, we measured the amplitude and rate of high frequency events identified using an anomaly detection algorithm (ADA) [40], which does not require application of an energy-based threshold. Second, we repeated the analysis using a conventional HFO detector based on root-mean-square (RMS) amplitude. For both detectors, we measured the rate and amplitude of detected events in SOZ and nSOZ electrodes and compared the classification accuracy based on these two features.

## 2. Methods

### 2.1 Patients and recordings

This study was approved by the Institutional Review Board of the University of California, Irvine. The iEEG recordings used in this study were collected from subjects with medically refractory epilepsy undergoing surgical evaluation at University of California, Irvine, Medical Center between April 2015 and December 2017. The selected subjects fulfilled the following inclusion criteria: (1) diagnosis of temporal lobe epilepsy; (2) intracranial monitoring resulted in the localization of the SOZ by epileptologists to one or more electrodes; (3) electrode locations were confirmed by co-registering pre- and post-implantation structural T1-weighted magnetic resonance imaging (MRI) scans; (4) availability of at least one full night of iEEG data (minimum six hours) containing no seizures; and (5) a minimum sampling frequency of 2 kHz. From 36 subjects who underwent intracranial monitoring to localize the SOZ for possible surgical resection, 11 adult subjects fit these criteria (five female, 38.2±16.9 years old; see Table 1). The data for 10 subjects had a 5 kHz sampling rate, and one subject had a 2 kHz sampling rate. The seizure onset times and SOZ were determined independently by experienced epileptologists (JJL and IS). The type of surgery performed, number of SOZ channels in the resected volume, and Engel outcome are provided for qualitative comparison (Table 1). Because the objective of this study is to measure HFO amplitude inside and outside the SOZ and assess the viability of this metric as a biomarker of the SOZ, we do not quantify the relationship between HFO amplitude and the surgery and outcome data. For the same reason, we also exclude non-SOZ channels exhibiting frequent epileptiform discharges and/or in regions of seizure spread; this provides the cleanest distinction between channels inside and outside the SOZ, while eliminating channels that require a more subjective assessment of whether or not they are abnormal and should be considered targets for resective surgery.

**Table 1:** Clinical data. Abbreviations: L = left, R = right, TLE = temporal lobe epilepsy, RNS = responsive neurostimulation. For the electrodes, AC = anterior cingulate, AH = anterior hippocampus, AM = amygdala, LES = left lesion, HH = head of hippocampus, IN = insula, OC = occipital, OF = orbitofrontal, P = parietal, PC = posterior cingulate, SM = supplementary motor area, TH = tail of hippocampus, TI = insula (transverse). For genders, F = female, M= male. For header, SOZ = seizure onset zone, nSOZ = non-seizure onset zone. * Resected volume was unavailable for this subject.

| Subject | Age (Gender) | Epilepsy Diagnosis | SOZ (# of bipolar pairs) | nSOZ (# of bipolar pairs) | # of clips analyzed | Surgery | SOZ channels resected | Outcome |
| --- | --- | --- | --- | --- | --- | --- | --- | --- |
| 1 | 23 (F) | LTLE | LTH (2) | LIN (2), RHH (1), RTH (2), LOF (2) | 8 | L temporal lobectomy | 1/2 | IIIA |
| 2 | 50 (M) | RTLE | RAH (3) | ROF (2), LIN (1), RIN (1) | 10 | R temporal lobectomy | * | IB |
| 3 | 69 (F) | RTLE | RAH (3) | ROF (2), RP (4), LP (2), LOF (1) | 12 | R temporal lobectomy | 2/3 | IIIA |
| 4 | 21 (M) | RTLE | RAM (1), RTH (1), RHH (2), LTH (2), LHH (2), LAM (2) | RTH (1), LTH (1), LHH (1), ROC (2) | 8 | R temporal lobectomy; ablation of hypothalamic hamartoma | 4/10 | IIB |
| 5 | 57 (F) | LTLE | LHH (2), LTH (2) | RAM (2), RTH (1), ROF (1), RIN (2), LAM (1), LTH (1), LAC (2) | 12 | L temporal lobectomy | 2/4 | IA |
| 6 | 34 (M) | Bilateral TLE | RAM (2), RHH (2), RTH (1), LAM (1), LTH (2) | ROF (1), RAC (2), RIN (1), RAM (1) | 13 | RNS | N/A | IIA |
| 7 | 22 (F) | LTLE | LHH (1) | LAC (2), LES (2), LSM (2), LIN (4), ROF (2), RIN (1) | 7 | RNS | N/A | IIIA |
| 8 | 41 (M) | LTLE | LAM (3), LHH (2), LTH (2) | LAM (1), LAC (3), LPC (2) | 10 | L temporal lobectomy | 2/3 | IIIA |
| 9 | 54<br>(F) | RTLE | RAM (1), RHH (3) | RAM (3), RTH (2),<br>RAC (2), LAM (4),<br>LHH (2), LTH (3),<br>LAC (1) | 10 | R temporal<br>lobectomy | 2/2 | IB |
| 10 | 23<br>(M) | RTLE | RHH (3), RTH (1) | RAC (2), RIN (3)<br>RTI (2), RHH (1),<br>LSM (2), LAC (2),<br>LOF (2), LAM (2),<br>LHH (1), LTH (2) | 17 | R temporal<br>lobectomy | 3/4 | ID |
| 11 | 27<br>(M) | Bilateral<br>TLE | RAM (3), RHH (3),<br>LHH (2), LTH (1) | RHH (2), RTH (2),<br>RAC (2), ROF (1),<br>RPC (2), LAM (3),<br>LHH (2), LTH (1),<br>LAC (4), LOF (2),<br>LPC (2) | 11 | RNS | N/A | IVB |

Fifty-five bipolar pairs of depth electrodes within the SOZ (5.0±2.9 channels per subject) and 119 pairs of depth electrodes in nSOZ (10.8±6.2 channels per subject) were selected for analysis. Each nSOZ channel was a bipolar pair in which both electrodes were clearly located within gray matter; channels located in white matter and those visually identified as having bad signal quality or continuous electrographic artifact were excluded (Supplementary S1). One subject had implanted electrocorticogram grids that were not involved in seizure onset, and they were excluded to ensure consistent electrode size across all subjects. Multiple three-minute clips of iEEG data were selected from each patient using the following procedure: (1) clips were selected from overnight iEEG recordings between 11 PM and 6 AM; (2) clips were at least one hour away from a seizure onset time; and (3) clips were at least 15 minutes apart, resulting in the selection of approximately three clips per hour. The number of clips for each subject ranged from 7 to 17. In total, 118 clips of iEEG (10.7±2.8 clips per subject or 32.1±8.4 minutes per subject) were analyzed in this study.

### 2.2 Anomaly detection algorithm (ADA)

ADA identifies high frequency events that stand out from the background signal without requiring assumptions about the appearance of those events [40]. The algorithm consists of data preprocessing, calculating the similarity between all 50 ms windows of data, and applying hierarchical clustering to the similarity matrix. In the original algorithm, all clusters except the largest (“background”) cluster were defined as anomalous [40]. Here, the anomalous clusters were selected using a threshold on the distance between the background cluster and the candidate anomalous cluster, where the threshold N is a multiplier for the standard deviation of intra-background cluster distances (see Supplementary S2). This change allowed us to evaluate the robustness of the results as the number of detected events varied. The default value was *N=1*, which resulted in detections that matched the original ADA. The algorithm was designed using an independent subset of data from three of the subjects, and it was then applied with fixed parameters to the entire dataset. The data used for algorithm development were not re-used to generate the results presented here.

### 2.3 RMS detector

The RMS detector was used to obtain the benchmark accuracy for classification of SOZ and nSOZ channels based on HFO rate, as it has been widely applied to a variety of intracranial electrode types across all regions of the brain [1,5,34,41,42]. Here, we implemented the algorithm with all parameters as defined in the original publication [43].

### 2.4 Characteristics of detected events

We detected events using ADA and RMS in all segments of data from all channels. Hereafter, we will refer to events detected by ADA as anomalous high frequency activity (aHFA) and events identified by the RMS detector as conventional HFOs (cHFO). All data analysis was done in MATLAB 2018b (MathWorks, MA) using custom-written code.

We measured two characteristics of aHFA and cHFO: (1) event rate, defined as the average number of detected events per minute per channel calculated for each segment of data, and (2) amplitude, defined for each event as the average of the upper amplitude envelope obtained via the Hilbert transform over the duration of the event. For each 3-minute segment of data, the average amplitude was calculated across all events detected within that segment. This resulted in one amplitude value per segment, which was used for segment-based comparisons (see Section 2.5). For channel-based comparisons (see Section 2.5), the segment-based amplitude values were averaged across all segments of data within a channel. This method treated all segments equally, rather than more heavily weighting the segments with more detections. If a segment or a channel contained zero detected events, it was assigned an amplitude of zero. For rate and amplitude, we also calculated the coefficient of variation (CV), defined for each channel as

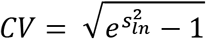

where s_ln_ is the sample standard deviation (SD) of the amplitude or rate across segments after a natural log transformation. In other words, the CV of each channel was calculated using the SD of the amplitudes or rates over all segments within that channel. Here, we used the CV to quantify the stability of the measurements across independent segments of data, in order to compare the values for amplitude and rate in SOZ and nSOZ channels.

### 2.5 Evaluation of biomarker performance

The ability to differentiate SOZ and nSOZ was evaluated via receiver operating characteristic (ROC) curves for event rate and amplitude for both detectors. Positives (P) were defined as the electrodes within the SOZ, as identified by epileptologists based on seizures captured via intracranial monitoring, and negatives (N) consisted of all brain regions outside the SOZ. The ROC curve was created by plotting the sensitivity, defined as TrueP/(TrueP+FalseN), against false positive rate (FPR), defined as FalseP/(FalseP+TrueN), for a range of amplitude and rate thresholds. This was done using two different methods: (1) A segment-based ROC curve was created by pooling all segments together and using the threshold to classify individual 3-minute segments, and (2) a channel-based ROC curve was created by applying the threshold to the average value for each channel, where the average was calculated across all iEEG segments in that channel. The mean values of rate and amplitude were used without any normalization to account for the different numbers of HFOs in each subject. Then the performance was assessed using the area under the ROC curve (AUC) and the sensitivity and FPR at the optimal cut-point. The AUC ranges from zero to one, with one indicating perfect performance. The optimal cut-point, which represents the best balance between sensitivity and FPR, was defined as the point that minimized the Euclidean distance between the ROC curve and the (0% FPR, 100% sensitivity) point at the upper left corner of the ROC plot.

We employed the Wilcoxon signed-rank test and Wilcoxon rank sum test to determine statistical significance for paired and unpaired samples, respectively, and the significance for all analyses was set at *p* < 0.05.

## 3. Results

### 3.1 The amplitude of high frequency events is higher in SOZ and is stable over time

Across SOZ channels, we detected a total of 6,208 aHFA and 14,008 cHFOs in 598 segments of iEEG data, where each segment contained three minutes of data from one channel. Across nSOZ channels, we detected 15,193 aHFA and 7,179 cHFOs in 1,336 segments of iEEG data.

Consistent with prior studies, the rate of cHFO within SOZ channels (8.2±4.2 per minute) was significantly higher than in nSOZ channels (1.9±2.7 per minute) (Figure 1A). However, the aHFA rate was significantly higher in nSOZ channels. Surprisingly, the amplitude of detected events was significantly higher in SOZ than nSOZ channels for both ADA and RMS detection (Figure 1B). Specifically, the amplitude of aHFA was 39.7±28.8 *μ*V in the SOZ and 7.2±8.7 *μ*V in the nSOZ, and the cHFO amplitude was 37.0±29.4 *μ*V in the SOZ and 6.4±8.1 *μ*V in the nSOZ.

**Figure 1:**
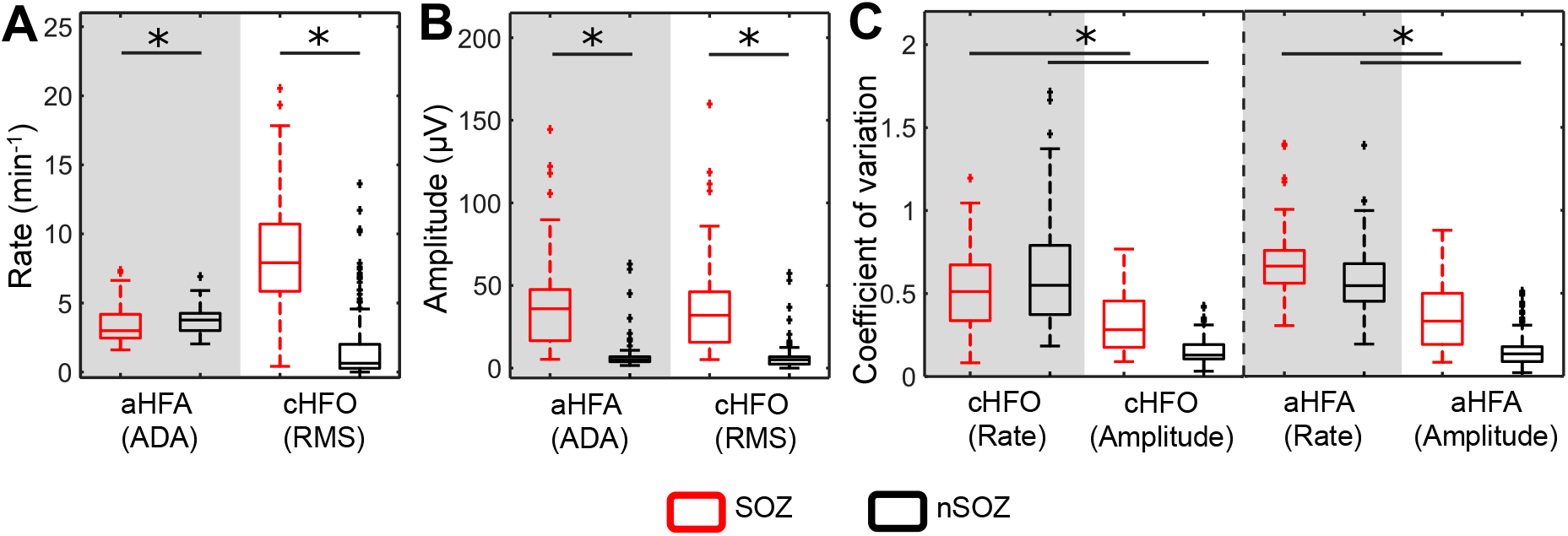
The conventional HFO (cHFO) rate, cHFO amplitude, and anomalous high frequency activity (aHFA) amplitude are higher in the seizure onset zone (SOZ), but the measurements of amplitude are more stable over time. (A) Rate and (B) amplitude of detected events from the anomaly detection algorithm (ADA) and root-mean-square (RMS) detectors. Each datapoint represents the mean value from one segment of iEEG. (C) Coefficient of variations (CVs) for rate and amplitude, calculated across all iEEG segments for each channel. All features are shown separately for SOZ (red) and non-SOZ (nSOZ) channels (black). *p-value < 0.001, Wilcoxon rank sum test.

The values of these metrics fluctuated over time, as previously reported by others [33]. However, we found that the amplitude measurement was more consistent across segments of data (i.e. more stable over time), evidenced by significantly lower CV values for the amplitude of aHFA and cHFO compared to rate (Figure 1C).

### 3.2 Amplitude enables more accurate classification of SOZ and nSOZ than rate for individual subjects

The individual subject results also reflect these differences in rate and amplitude; this is crucial, as HFO analysis will typically be done on single subjects if used clinically for SOZ localization. The heatmaps in Figure 2 depict the differences between SOZ and nSOZ channels for all iEEG segments for each individual subject. Visually, both the amplitude and rate of detected events were higher in the SOZ than the nSOZ channels, as expected; however, the amplitude measurements typically exhibited a greater difference between SOZ and nSOZ channels. In particular, the amplitude values in nSOZ channels were lower relative to the SOZ values and were more consistent over time.

**Figure 2:**
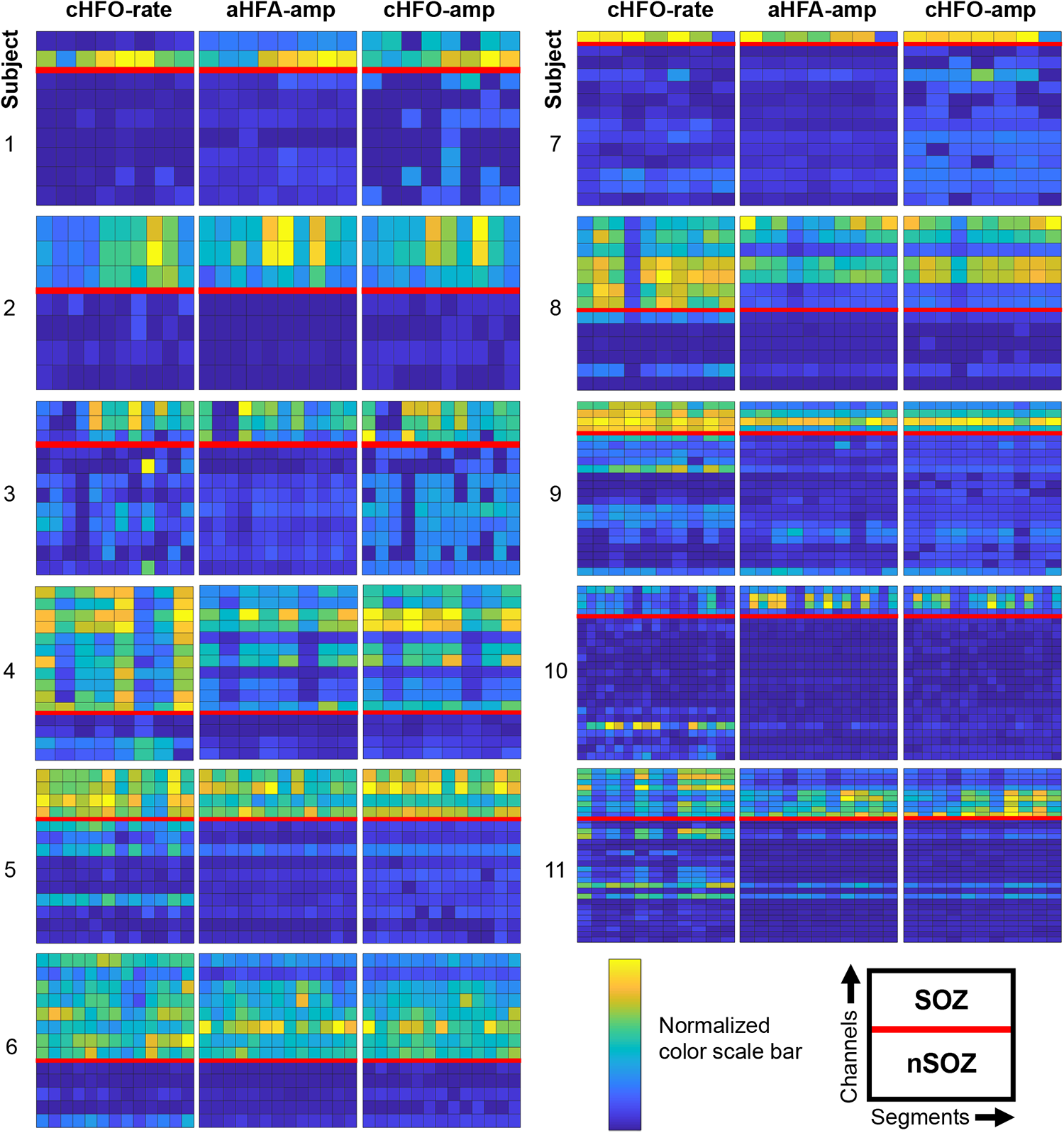
Amplitude exhibits greater differences between seizure onset zone (SOZ) and non-SOZ (nSOZ) channels than conventional HFO (cHFO) rate. Heatmaps illustrate the cHFO rate (left sub-panel), anomalous high frequency activity (aHFA) amplitude (center sub-panel) and cHFO amplitude (right sub-panel) for all channels and all iEEG segments. Each set of three sub-panels shows data from one subject. In each sub-panel, each column represents data from one 3-minute iEEG segment for all channels. Each row shows the data from a single channel, with the channels divided into SOZ (above red line) and nSOZ (below red line). The color represents the mean rate or amplitude of the detected events during one 3-minute segment of iEEG, normalized by dividing by the maximum value within each individual subject.

To assess the performance of all three metrics as potential SOZ biomarkers, we calculated the segment-based ROC curves for all individual subjects. The ROC curves for amplitude were generally closer to the upper left-hand corner, suggesting that amplitude provided better separation between SOZ and nSOZ channels than cHFO rate (Figure 3A-C). To statistically compare the performance of the three metrics, we used a Wilcoxon signed-rank test to assess the differences in AUC, as well as sensitivity and FPR at the optimal cut-point of individual subjects (Figure 3D). The AUCs of the aHFA amplitude and cHFO amplitude (mean values of 0.960 and 0.948, respectively) were significantly higher than the AUC for the cHFO rate (0.905). Furthermore, the FPR values at the optimal cut-points of the aHFA amplitude (mean of 6.1%) and cHFO amplitude (6.2%) were significantly lower than for cHFO rate (13.7%). For the sensitivity, the aHFA amplitude (mean of 93.7%) and cHFO amplitude (mean of 94.1%) were significantly higher than the cHFO rate (84.9%) at a *p*-value of 0.05 (Figure 3D). In comparing the ADA and RMS detectors, the aHFA amplitude had a higher mean AUC and lower mean FPR than cHFO amplitude, but these differences were not statistically significant.

**Figure 3:**
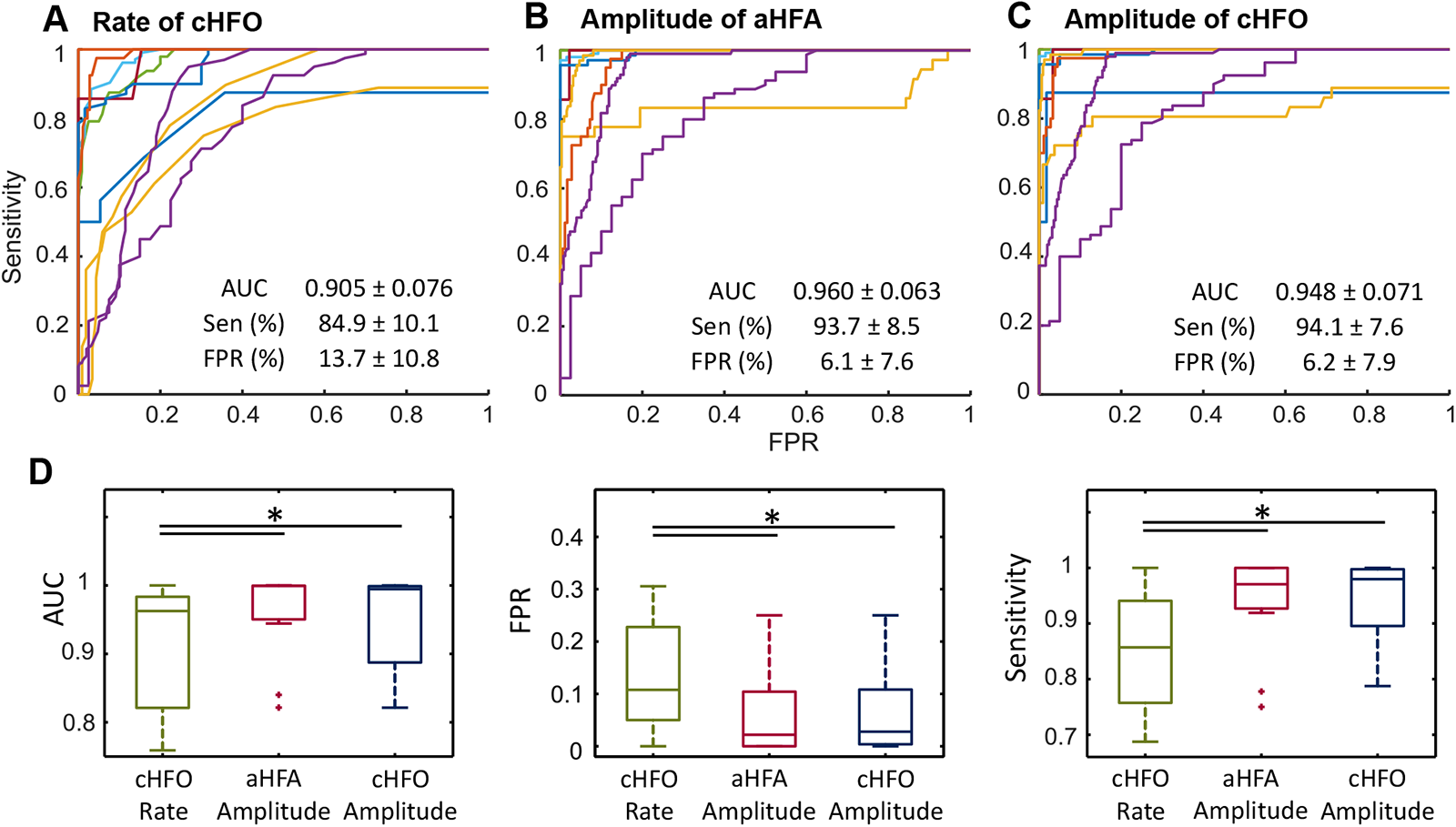
For classification of seizure onset zone (SOZ) and non-SOZ (nSOZ) channels in individual subjects, amplitude provides better performance than conventional HFO (cHFO) rate. Receiver operating characteristic (ROC) curves are shown for (A) cHFO rate, (B) anomalous high frequency activity (aHFA) amplitude, and (C) cHFO amplitude for classification of SOZ and nSOZ channels. The ROC curves were calculated using the segment-based scheme, and each line in the subfigures represents an individual subject. (D) Boxplots of area under the curve (AUC), false positive rate (FPR), and sensitivity at the optimal cut-point for all 11 subjects computed from the ROC curves in (A-C). Results are shown for cHFO rate (green), aHFA amplitude (red), and cHFO amplitude (blue). The amplitude metrics have significantly higher AUC and lower FPR than the cHFO rate, and the cHFO amplitude has significantly higher sensitivity than cHFO rate. *p-value < 0.05, Wilcoxon signed-rank test.

### 3.3 Amplitude enables more accurate classification of SOZ and nSOZ than rate at the group level

The overall difference in performance between the amplitude and rate metrics can be observed by generating the segment-based (Figure 4A) and channel-based (Figure 4B) ROC curves for all patients pooled together. Both ROC curves gave comparable results. The segment-based ROC curves indicated similar performance between the amplitude of aHFA and cHFO (AUC: 0.946 and 0.942, respectively), and both outperformed the rate of cHFO (AUC: 0.880) (Figure 4A). Similarly, for the channel-based method, the AUC values were 0.956 for aHFA amplitude, 0.952 for cHFO amplitude, and 0.911 for cHFO rate (Figure 4B). The segment-based method demonstrates the performance across measurements from different points in time, and the channel-based method matches the potential clinical use of the metric, where individual channels would be classified as SOZ or nSOZ during surgical planning. Both approaches indicate the robustness of the amplitude metrics when using a common cut-point across all subjects, as opposed to the individualized cut-points used in Figure 3.

**Figure 4:**
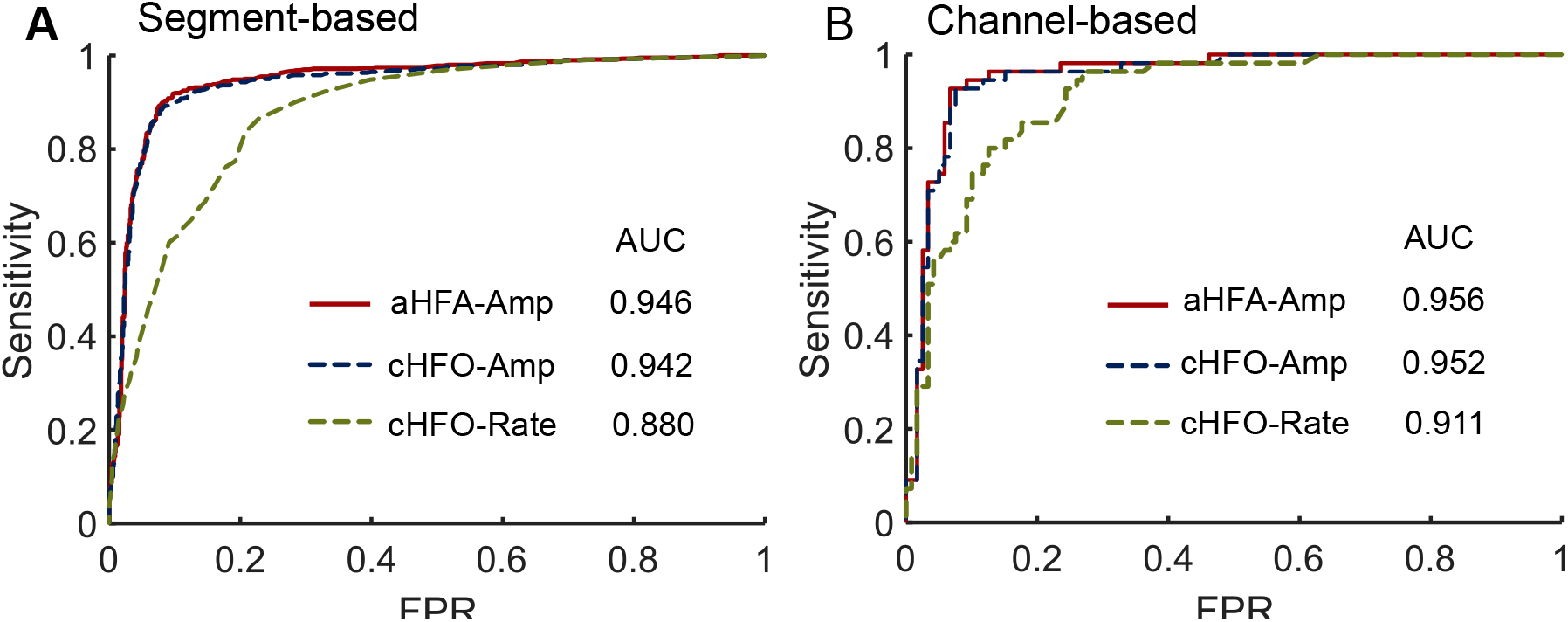
Amplitude provides superior classification of seizure onset zone (SOZ) and non-SOZ (nSOZ) channels with data pooled across all subjects. Receiver operating characteristic (ROC) curves are shown for conventional HFO (cHFO) rate (green), anomalous high frequency activity (aHFA) amplitude (red), and cHFO amplitude (blue). (A) ROC curves calculated using the segment-based method. (B) ROC curves calculated using the channel-based method. The amplitude metrics result in higher area under the curve (AUC), higher optimal sensitivity, and lower optimal false positive rate (FPR) using both evaluation methods.

### 3.4 The amplitude metric is more robust than rate to changes in detection sensitivity

In addition to the advantage in the measurement consistency (Figure 1C), we found that the amplitude was more robust to changes in the detection parameters than the rate. We varied the detection threshold for each algorithm (the value of *N* in the separation threshold for ADA and the number of standard deviations used for the RMS threshold in the RMS detector), and we measured the resulting AUC values after classification of SOZ and nSOZ using the segment-based approach (Figure 5). The AUC values for the aHFA rate and cHFO rate were highly dependent on the threshold, peaking at a mid-range value. For example, varying the value of *N* in ADA caused the AUC for aHFA rate to increase from 0.395 (*N*=1) to a max of 0.848 (*N*=7). Similarly, for cHFO rate, the AUC increased from 0.565 (*N*=1) to a max of 0.901 (*N*=9). This shows that optimization of the threshold is critical to obtaining high accuracy when using the rate of events as a metric for classification. In contrast, the AUC values computed using the aHFA amplitude (range of 0.817 (*N*=11) to 0.946 (*N*=1)) and cHFO amplitude (range of 0.783 (*N*=17) to 0.928 (*N*=1)) were relatively robust to alteration of the threshold. Generally, for amplitude, the best results were obtained for the lowest thresholds.

**Figure 5:**
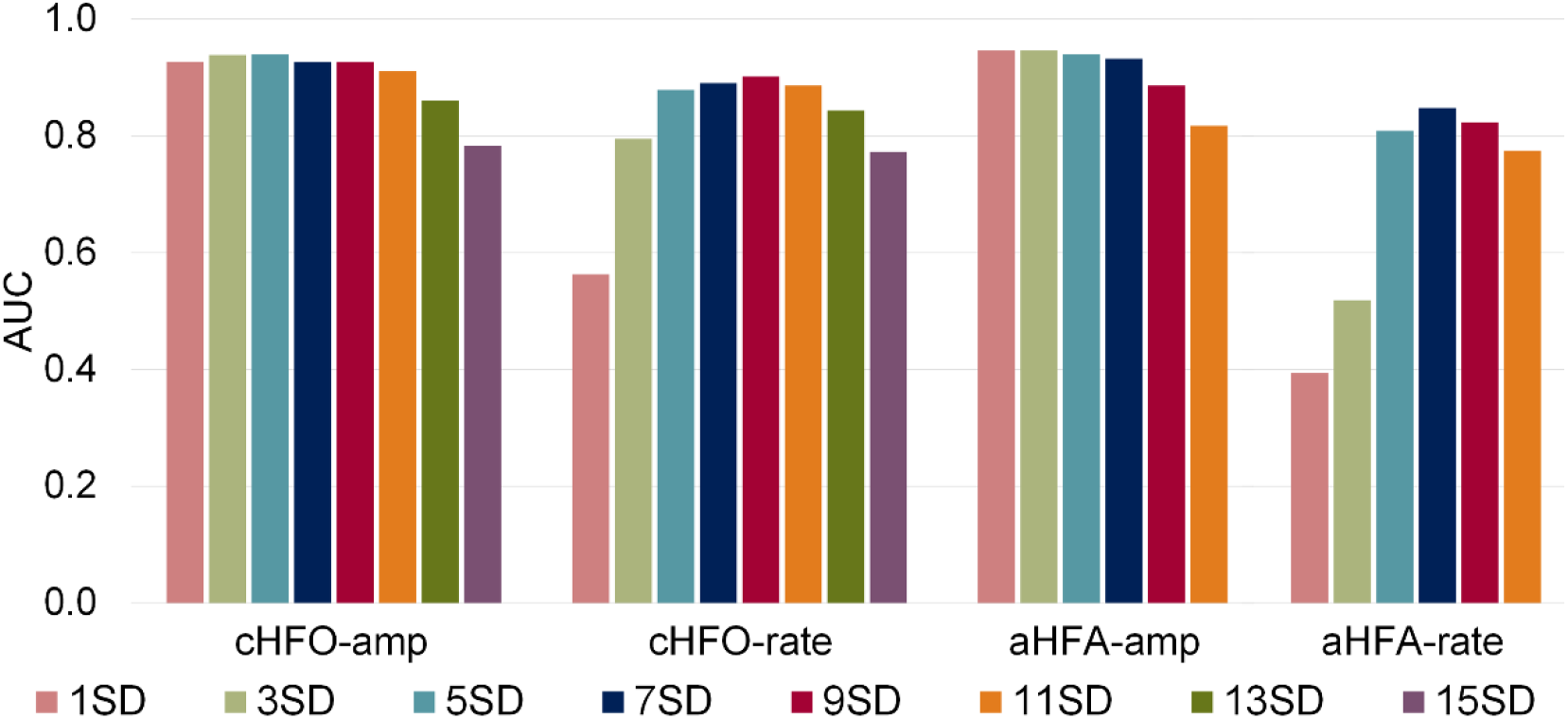
The amplitude metric is robust to changes in the detector threshold. For anomaly detection algorithm (ADA), the number of standard deviations (SD) above the mean in the separation threshold was varied from N=1 to N=11. For the root-mean-square (RMS) detector, the threshold applied to the RMS amplitude was varied from one to 15 standard deviations above the mean. Area under the curve (AUC) values are shown for the amplitude and rate of anomalous high frequency activity (aHFA) and conventional HFO (cHFO) using the segment-based calculation. For N<13, the AUC for the amplitude metric is consistently higher than 0.8, with the highest AUC values occurring at the lowest values of N.

## 4. Discussion

Here, we present evidence that the HFO amplitude is a candidate biomarker of the SOZ with high localization accuracy, measurement stability, and robustness to changes in detection parameters. Consistent with prior studies, we found that the rate of cHFO can be used to classify SOZ and nSOZ electrodes (average AUC of 0.905, with 84.9% sensitivity and 13.7% FPR at the optimal cut-point). However, in a head-to-head comparison, using aHFA amplitude for classification led to a 6% increase in AUC, 11% increase in sensitivity, and 55% decrease in FPR relative to the cHFO rate. The performance using cHFO amplitude was similar to that of aHFA amplitude. Moreover, we found that the amplitude measurements were more consistent over time, as indicated by lower CV values, and the classification performance using amplitude was robust to changes in the detection threshold. Overall, this suggests that HFO amplitude may improve SOZ localization during epilepsy surgical planning, whether used alone or in conjunction with measurements of HFO rate.

Previous literature supports the idea that HFO amplitude is a potential marker of the SOZ [14,22,23,26,27,37,39]. One study suggested that an amplitude difference in the fast ripple band is specific to patients with good surgical outcome [27]. Moreover, pathological HFOs show higher mean spectral amplitude than induced physiological HFOs [37]. However, the reported amplitude distributions for SOZ and nSOZ channels were often overlapping [22,37], and classification using HFO amplitude was generally inferior to rate [22,37,44]. We hypothesized that this was due to the choice of automated detection algorithm and the selection of detection parameters. Consistent with this hypothesis, our results showed that as the threshold increased, the amplitude difference between SOZ and nSOZ decreased (Figure 5). All prior measurements of HFO amplitude used some form of energy threshold for detection, with some using highly specific techniques based on the time-frequency decomposition [14,23]. One study required that all HFOs, in both SOZ and nSOZ, have an amplitude greater than 10 μV [26], which would have excluded most of the events we detected in nSOZ electrodes (Figure 1B). Similarly, visual detection or confirmation of HFOs [22,23], the selection of detection parameters based on visual markings [14,27], or visually selecting a subset of channels for analysis [37] can bias detection toward the highest amplitude events in both SOZ and nSOZ, thus obscuring the difference between them. It is noteworthy that the study reporting the largest differences in amplitude between SOZ and nSOZ used a highly sensitive detector [39].

While the characteristics of HFO rate and amplitude are measured in different ways, they are not independent variables. For example, when using the RMS detector, both will be influenced by the detection threshold. As the threshold is increased, the number of detected HFOs (and thus, the rate) will decrease, but the amplitude of events will increase. In many cases, if HFOs in the SOZ have a higher amplitude than in nSOZ channels, then a correctly selected amplitude threshold should also result in good classification based on HFO rate. This is supported by the results in Figure 5, where the best overall performance using HFO rate (at a threshold of 9 SD) is similar to the performance using HFO amplitude with a lower threshold.

On the other hand, there could be scenarios that favor amplitude over rate. The threshold for detection is typically based on a number of standard deviations above the mean, and it is tailored to the activity in each channel. For example, 3 SD above the mean in the SOZ could be 10 μV, while 3 SD in nSOZ may only be 4 μV. Even an nSOZ channel that is purely noise will have some HFO-like events that exceed this threshold by chance, which can be confirmed by performing detection on pink noise filtered in the ripple band. However, if the SOZ channel contains prominent, high amplitude HFOs, this can skew the amplitude distribution and make it non-Gaussian. Overall, this will increase the threshold, thereby decreasing the number of chance detections, while true HFOs will be still detected. Therefore, in this case, both channels could have approximately the same rate, although the detections in the SOZ would demonstrate higher amplitude.

The ADA and RMS detectors identified an overlapping set of events in the high frequency iEEG, with subsets of events detected only by ADA and only by RMS [40]. Generally, the amount of overlap between aHFA and cHFO increased as the threshold for each detection algorithm increased. At a threshold of *N*=1, 82% of the aHFA in the SOZ were also detected by the RMS detector with the standard threshold of 5 SD. This percentage increased to 91% for *N*=5. This indicates that ADA detects events in the SOZ that fit the conventional, empirical definition of an HFO. In nSOZ channels, the percent overlap was lower, increasing from 21% to 50% for *N*=1 to *N*=5. This is because ADA detected many low and intermediate amplitude events which did not exceed the RMS amplitude threshold. However, this difference between the two detectors is expected. The purpose of the anomaly detector is to identify events that stand out from the background, without prescribing the characteristics of these events. For example, in developing this detector, we looked for parameters that assigned a vast majority of the signal to the background cluster, with relatively rare detection of unique events. We did not require the detected events to have specific amplitude or oscillatory features, as the goal was not to mimic visual HFO detection.

Given this, it was surprising that both ADA and the RMS detector provided estimates of HFO amplitude that could be used to classify SOZ and nSOZ channels. The differences in performance between the two detectors were not statistically significant. This suggests that existing detection algorithms, if sufficiently sensitive, can be used to measure HFO amplitude instead of rate. Because both ADA and the RMS detector are susceptible to false positive detections (where “false positive” is defined relative to visually-identified events), it is likely that some of the events used to estimate amplitude would not fit the traditional empirical definition of an HFO, especially in nSOZ channels. However, here we define cHFO and aHFA as any events that fit the criteria of the detection algorithms, regardless of their visual appearance, so these events are not false positives in the context of this study. We found that, because the non-traditional events detected in nSOZ channels tended to have very low amplitude, they were distinct from the high-amplitude events seen in SOZ regions. In contrast, when measuring the HFO rate using ADA, detection of these non-traditional events led to a high rate in nSOZ, thus obscuring the difference between SOZ and nSOZ regions.

In the long term, new detection algorithms and strategies for parameter optimization should be developed specifically for amplitude. For example, ADA offers several potential advantages over other algorithms. It does not require any training or parameter optimization, it is fully unsupervised, and it does not require prior assumptions about the shape or amplitude of the high frequency events. Figure 4 shows a trend of better performance for ADA amplitude versus RMS amplitude; analysis of a larger cohort of subjects and investigation into the causes of outliers can determine whether this difference is real. One disadvantage of ADA is that the approximate processing time to differentiate unique events for three minutes of data from a single channel of iEEG was approximately ~4-5 minutes using a desktop PC (CPU: i7-4790k).

In addition to higher classification accuracy for HFO amplitude compared to HFO rate, we found that the performance using amplitude was more robust to changes in detection parameters. For both ADA and the RMS detector, we showed that high AUC values for amplitude could be achieved with a wide range of detection thresholds (Figure 5). Therefore, ADA can be used as originally designed [40], without the additional separation threshold parameter. In the ADA clustering and classification process, we used a maximum of seven clusters; however, we tested a range of seven to fifteen maximum clusters and found almost no difference in the amplitude AUC (Supplementary S3).

There were several limitations to our study. We analyzed data from 11 subjects, with approximately 30 minutes of iEEG per patient; however, this is a relatively small number of subjects, considering the diversity of epilepsy syndromes. Moving forward, it will be necessary to obtain a larger cohort of subjects and analyze a larger percentage of electrodes, while accounting for the exact resected brain region and long-term surgical outcome. Here, we analyzed only electrodes that were clearly localized to gray matter. SOZ electrodes were defined based on the clinical assessment of the patient’s seizures. The nSOZ group included all other electrodes, except those with frequent epileptiform discharges and/or in regions of seizure spread. These strict criteria for channel selection likely strengthened our SOZ localization results for both HFO amplitude and rate, especially relative to studies in which all electrodes were included in the analysis. Because the ability to identify potential surgical targets among all implanted electrodes is critical to the success of epilepsy surgery, future studies will incorporate all electrodes into the analysis, as well as addressing the other limitations. The long-term goal of this work is to identify robust, accurate biomarkers of the epileptogenic zone to aid surgical planning. Ultimately, this will increase the percentage of patients with refractory epilepsy who are seizure free after surgery and may also increase the number of patients who are offered surgery as a treatment option.

## Acknowledgements

This research was financially supported by an American Epilepsy Society Junior Investigator Research Award and a Royal Thai Government Fellowship.

