## Supplementary for "Amplitude of high frequency oscillations as a biomarker of the seizure onset zone"

**Supplementary S1: Electrode information for individual subjects**

Counts are shown for all depth electrodes implanted in the 11 subjects included in this study. The total number of channels is given, and the subsets of gray matter channels in the SOZ, region of seizure spread, and nSOZ are listed in columns 3-5. The numbers of channels rejected based on localization are listed in columns 6-8. Note that each channel in this table represents one electrode, while bipolar pairs were used in the analysis. The paired electrodes needed to be adjacent and in gray matter in order to create a bipolar pair, so the number of analyzed channels (in Table 1) is lower than the number listed here.

| Subject | Total number of channels | SOZ Channels | Spreading Channels | nSOZ Channels | White Matter | Out of the Brain | Unclear/On border* |
|---|---|---|---|---|---|---|---|
| 1 | 100 | 4 | 8 | 30 | 10 | 33 | 15 |
| 2 | 80 | 4 | 15 | 19 | 22 | 5 | 15 |
| 3 | 60 | 4 | 13 | 18 | 10 | 7 | 8 |
| 4 | 56 | 12 | 0 | 14 | 19 | 3 | 8 |
| 5 | 122 | 6 | 14 | 38 | 22 | 32 | 10 |
| 6 | 110 | 13 | 0 | 32 | 37 | 7 | 21 |
| 7 | 148 | 2 | 12 | 53 | 28 | 37 | 16 |
| 8 | 130 | 10 | 80 | 18 | 17 | 0 | 17 |
| 9 | 100 | 5 | 6 | 54 | 24 | 6 | 5 |
| 10 | 160 | 6 | 9 | 67 | 35 | 10 | 33 |
| 11 | 120 | 11 | 0 | 62 | 30 | 4 | 13 |

*Unclear/On border: Marked as more than one region, the location was not clearly marked, marked as "edge"

**Supplementary S2: Anomaly detection algorithm**

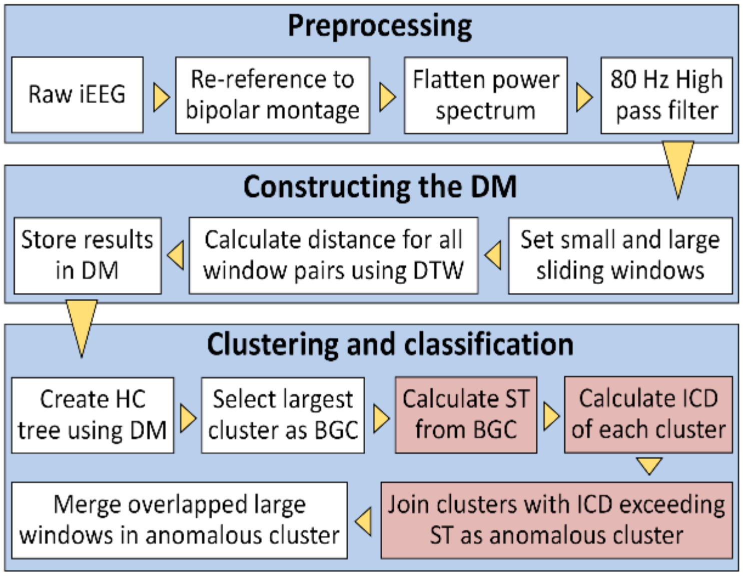


Flow chart for modified ADA. The algorithm consists of three main parts: preprocessing, constructing the DM, and clustering and classification. The white boxes represent the major processing steps for ADA, and the red boxes show the steps added to the original algorithm. Arrowheads show the flow of the algorithm. Abbreviations are DM: distance matrix, DTW: dynamic time warping, HC: hierarchical cluster, BGC: background cluster, ST: separation threshold, ICD: inter-cluster distance.

To identify anomalous events, we started by selecting the main background cluster (BGC), defined as the single cluster with the largest number of members. Here each member is one 50 ms window of data. In the original algorithm, all clusters except the BGC were defined as anomalous.[40] Here, the anomalous clusters were selected by comparing the inter-cluster distance (ICD) to a separation threshold calculated from the pairwise distances of all members within the BGC:

$$\text{Separation threshold} = \bar{d} + N\sqrt{\frac{1}{n}\sum_{i=1}^{n}\sum_{j=1}^{n}(d_{ij} - \bar{d})^2}$$

where $d_{ij}$ is the pairwise distance between windows i and j in the BGC; $\bar{d}$ is the mean of all pairwise distances $d_{ij}$; *n* is the number of members of BGC; and *N* is the number of standard deviations (SD). The ICD(*k*) is the mean distance between the *k*th cluster, $C_k$, and the BGC:

$$ICD(k) = \sum_{i \in C_k}\sum_{j \in BGC} d_{ij}/(m \times n),$$

where $d_{ij}$ is the distance between the *i*th member of $C_k$ and the *j*th member of the BGC, and *m* and *n* are the number of members in $C_k$ and the BGC, respectively. Any cluster with an ICD exceeding the separation threshold was marked as an anomalous cluster. Within the anomalous clusters, any 50 ms windows within one channel that overlapped in time were merged into a single event. We used a default value of *N=1* which resulted in detections that matched the original ADA.

**Supplementary S3**: **Classification results are robust to changes in the maximum number of clusters in the ADA clustering and classification process**.

For ADA, the separation threshold was set to its default value (*N*=1). The maximum number of clusters was varied from seven (light blue) to fifteen (dark blue). Results show the AUC calculated from the ROC curves of the amplitude and rate of aHFA using the segment-based and channel-based methods. The performance is consistently high for the amplitude metric. On the contrary, the results for rate are variable and depend on the choice of the maximum number of clusters; however, the effect is far less than the choice of threshold. This also confirms the finding that the change in the population of detected events, which can be seen from the variation of AUCs of the rate, did not affect the amplitude as a metric of SOZ separation.

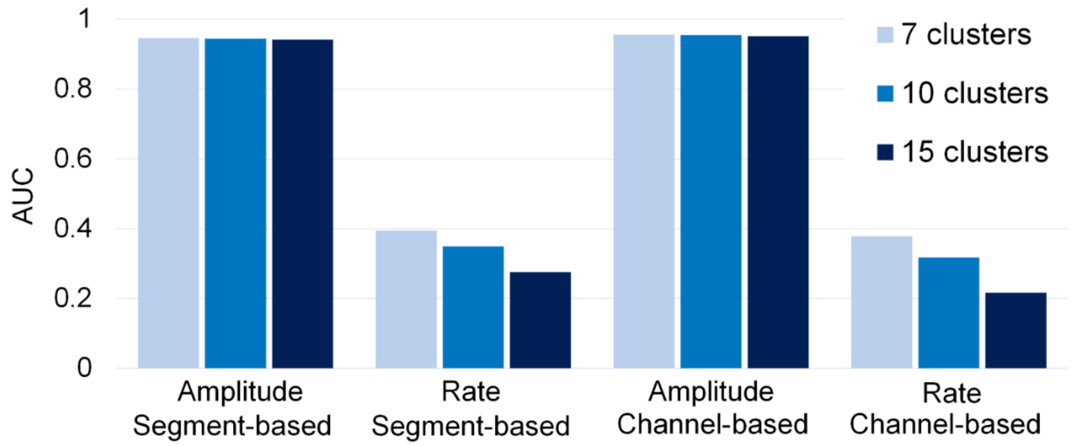
